# Forseti: A mechanistic and predictive model of the splicing status of scRNA-seq reads

**DOI:** 10.1101/2024.02.01.577813

**Authors:** Dongze He, Yuan Gao, Spencer Skylar Chan, Natalia Quintana-Parrilla, Rob Patro

## Abstract

**Motivation:** Short-read single-cell RNA-sequencing (scRNA-seq) has been used to study cellular heterogeneity, cellular fate, and transcriptional dynamics. Modeling splicing dynamics in scRNA-seq data is challenging, with inherent difficulty in even the seemingly straightforward task of elucidating the splicing status of the molecules from which sequenced fragments are drawn. This difficulty arises, in part, from the limited read length and positional biases, which substantially reduce the specificity of the sequenced fragments. As a result, the splicing status of many reads in scRNA-seq is ambiguous because of a lack of definitive evidence. We are therefore in need of methods that can recover the splicing status of ambiguous reads which, in turn, can lead to more accuracy and confidence in downstream analyses.

**Results:** We develop `Forseti`, a predictive model to probabilistically assign a splicing status to scRNA-seq reads. Our model has two key components. First, we train a binding affinity model to assign a probability that a given transcriptomic site is used in fragment generation. Second, we fit a robust fragment length distribution model that generalizes well across datasets deriving from different species and tissue types. `Forseti` combines these two trained models to predict the splicing status of the molecule of origin of reads by scoring putative fragments that associate each alignment of sequenced reads with proximate potential priming sites. Using both simulated and experimental data, we show that our model can precisely predict the splicing status of reads and identify the true gene origin of multi-gene mapped reads.

**Availability:** `Forseti` and the code used for producing the results are available at https://github.com/COMBINE-lab/forseti under a BSD 3-clause license.

## Introduction

Single-cell RNA-sequencing (scRNA-seq) technology has revolutionized our understanding of cellular heterogeneity and differentiation dynamics (Stark et al., 2019), and short-read, 3^1^, tagged-end technologies have dominated contemporary data generation. In the most popular scRNA-seq protocols, poly(T) primers are used to capture the polyA tail of polyadenylated RNAs. However, recent studies have shown that intronic reads usually account for ∼ 20% to ∼ 40% of the total gene count (i.e. distinct number of unique molecular identifiers UMIs) in scRNA-seq data (He et al., 2023; 10x Genomics, 2021), suggesting that, in addition to the polyA tails, the internal adenine-single nucleotide repeats (A-SNR or polyA) — predominantly on unspliced transcripts — are also frequently primed by oligo(dT) primers in scRNA-seq to generate sequenced scRNA-seq reads (Nam et al., 2002; Svoboda et al., 2022; 10x Genomics, 2021). Prior work has also shown that the information captured from unspliced transcripts can offer unprecedented insights into single-cell biology from a brand new perspective Pool et al. (2023); Chamberlin et al. (2022); 10x Genomics (2022b); Gorin et al. (2023). For example, single-cell RNA velocity (La Manno et al., 2018) infers the cellular differentiation dynamics by proposing and performing inference in a model that uses the spliced and unspliced scRNA-seq reads separately to infer the transcriptional dynamics of the underlying genes and cells. This innovation extended the horizon of single-cell data analysis, and has inspired a plethora of subsequent works (Bergen et al., 2020; Li et al., 2023).

Although the community has been paying increasing attention to the development and use of novel algorithms that utilize information captured from unspliced transcripts, a fundamental problem has yet to be solved. Specifically, the accurate identification of the splicing status of scRNA-seq reads remains a difficult challenge (He et al., 2023, 2022; Eldjárn Hjörleifsson et al., 2022a). Currently, the standard strategy of assigning splicing status to reads follows the heuristics introduced in La Manno et al. (2018), in which fully and partially intronic reads are classified as unspliced reads, and reads that are only compatible with exonic regions are classified as spliced reads. However, this strategy implies a strong preference for classifying reads as spliced. This is because reads that are entirely contained within the body of an exon, which can originate from either spliced or unspliced transcripts, are all classified as spliced reads. An alternative strategy is to assign an ambiguous splicing status to these exonic reads, indicating that the actual splicing status of the reads is undetermined (Eldjárn Hjörleifsson et al., 2022a; He et al., 2023). However, as a tremendous fraction of scRNA-seq reads are exonic (10x Genomics, 2021), applying this strategy leads to, on average, over half of the gene counts (UMIs) being assigned as ambiguous (∼ 46% to ∼ 62% for the eight datasets processed in He et al. (2023), with a mean of ∼ 53%).

In general, in the current work, we refer to “unspliced” molecules in the understanding that they might be undergoing splicing and hence, may be partially spliced at the point when the cell was lysed. Further, having an “ambiguous” splicing status indicates that, for a read, the splicing status of its transcript origin, which in reality will be either spliced or unspliced, cannot be determined because of a lack of definitive evidence.

In this work, we introduce `Forseti`, the first probabilistic model of which we are aware for resolving the splicing status for exonic scRNA-seq reads. Our model does this by taking advantage of the technical details of the underlying scRNA-seq technologies. Specifically, our model is based on the fact that the expected priming sites of oligo(dT) primers are A-SNR, and the synthesized cDNAs are fragmented with a length preference of ∼ 300 to ∼ 400 base pairs^*^, and follow a distribution that is well-concentrated about the mean. By utilizing the fragment length distribution computed from publicly available, paired-end 3^1^ scRNA-seq datasets, and a multilayer perceptron model trained using the priming sites obtained from these datasets, `Forseti` obtains an Area Under the Receiver Operating Characteristic Curve (ROC AUC) score of 93.5% on simulated data, and over 90% on two experimental datasets.

We believe that this work represents a substantial and important step in improving the specificity and accuracy of gene quantification, and specifically splicing status determination, from scRNA-seq data. After describing and evaluating our model to demonstrate its utility, we discuss how the predictions of such a model might immediately aid in improving the downstream processing of scRNA-seq data, but also how the probabilistic allocations produced by our model may, themselves, help to enable more accurate and robust processing, as well as how the model might be extended and enhanced in the future.

## Methods

### Data Description

In this study, we collected 10 publicly available scRNA-seq datasets using 10X Chromium v2 and v3 solutions for training and testing our model. These datasets consist of both nucleus samples and cell samples from human and mouse across multiple tissue types (table 1). All these datasets are generated using the alternative sequencing formats (10x Genomics, 2022a), by which the technical read (Read1) is sequenced with the same number of cycles as the biological read (Read2), and therefore, contains the 5′ end of the read’s cDNA insert (10x Genomics, 2022a). Compared with the standard sequencing formats, in which reads1 contains only the technical barcode sequences, the reads from the datasets used in this work can be processed as “paired-end” datasets, so as to be used to calculate the cDNA fragment length in scRNA-seq (section 2.3). We processed all human datasets against the GRCh38 version 2020-A genome build and all mouse datasets against the mm10 version 2020-A genome build. Both genome builds were downloaded from the 10X Genomics website ^†^. The corresponding gene annotations were downloaded together with the genome builds and were used to make the augmented gene annotation set. We utilized eight datasets for model training and two datasets for model evaluation. The training and test datasets were randomly selected and are marked in the *train/test* column in table 1.

**Table 1.** Details of the ten datasets used in this work. the *Chemistry* column records the version of the 10x Chromium 3′ assay used for the corresponding datasets. The *train/test* column represents the label of the random split of the datasets used for training and testing `Forseti`. All datasets are publicly available and can be downloaded from NCBI GEO at https://www.ncbi.nlm.nih.gov/geo/.

| # | GSE ID | Species | Cell type | Chemistry | Read length | Cells or Nuclei | train/test |
| --- | --- | --- | --- | --- | --- | --- | --- |
| 1 | GSE144136 | Human | Brain | v2 | 100 | Nuclei | train |
| 2 | GSE148504 | Human | Cardiomyocytes | v3 | 150 | Cells | train |
| 3 | GSE122743 | Human | MCF7 | v2 | 75 | Cells | train |
| 4 | GSE125970 | Human | Ileum/Colon/Rectum | v2 | 150 | Cells | train |
| 5 | GSE130636 | Human | Retina | v3 | 150 | Cells | train |
| 6 | GSE131736 | Human | Retina | v2 | 75 | Cells | train |
| 7 | GSE134520 | Human | Stomach | v2 | 150 | Cells | train |
| 8 | GSE135922 | Human | RPE/Choroid | v3 | 151 | Cells | train |
| 9 | GSE122357 | Mouse | Brain | v2 | 151 | Cells | test |
| 10 | GSE125188 | Human | Liver/Blood/Spleen | v2 | 150 | Cells | test |

### Augmented gene annotations

In this work, we built an augmented gene annotation set for mouse and human, which we denote as transcript-level *spliced+unspliced* references, or *spliceu* in short, to obtain the read compatibility to both spliced and unspliced transcripts. The *spliceu* references were generated by adding the unspliced version of each multi-exon transcript to the standard 10X gene annotations (section 2.1) in an `R` (version 4.3.2) environment, with the help of the `BSgenome` (Pagès, 2023), `GenomicRanges` and `GenomicFeatures` (Lawrence et al., 2013), `eisaR` (Soneson et al., 2021), `Biostrings` (Pagès et al., 2023), and `rtracklayer` (Lawrence et al., 2009) packages. The unspliced version of a transcript is a contiguous genomic interval from the 5′-most exonic locus to the 3′-most exonic locus of the transcript. The single-exon transcripts (mainly non-coding RNAs) were not augmented in *spliceu* reference, because they do not have introns to be spliced. We first loaded the 10X genome build and the exon by transcript information using the `readDNAStringSet` function from `Biostrings` and `getFeatureRanges` from `eisaR`. The exon annotations were then written as a `BED` file using `export.bed` function from `rtracklayer`. Unspliced transcript annotations were defined as the range of transcripts. The sequence of spliced and unspliced transcripts were then obtained according to the genome and the spliced and unspliced transcript annotations using `extractTranscriptSeqs` from `GenomicFeatures`. With the transcript sequences in hand, we also found the A-SNR on each transcript using the `vmatchPattern` function from `Biostrings`. Here a A-SNR is defined as an adenine-single nucleotide repeat (aSNR) of length 6 or greater without mismatch. The A-SNR information will be used to assist in simulation in later steps.

### Read preprocessing

The composition of a cDNA fragment in the scRNA-seq technologies we consider here has been clearly explained in 10x Genomics (2022a) and Chen et al. (2023). Briefly, in scRNA-seq, from the 5′ end to the 3′ end of a first strand cDNA, apart from all PCR and sequencing primers, consists of a cellular barcode (CB) of length 16 base pairs, a unique molecule identifier (UMI) of length 10 or 12 base pairs depending on the chemistry, a poly(dT) sequence of length 32 bases corresponding to the oligo(dT) primer, and a cDNA insert of a various length but with a preference of 190 to 290 base pairs ^*^ (without technical oligonucleotides, such as CB, UMI and primers). When following the alternative sequencing format (10x Genomics, 2022a), the corresponding read 1 consists of CB, UMI, a poly(T) sequence corresponding to the priming window, and the 5′ end of the cDNA insert. The Read2 sequence consists of the 3′ end of the cDNA insert (i.e. the standard “biological” read2). Throughout this work, we define the CB, UMI and poly(T) sequence in read 1 as technical read 1, and the cDNA insert in read 1 as biological read 1. In the following text, we discuss the procedure used for processing the sequencing reads of each selected dataset.

In order to align the biological read 1 and read2 together, we first break read 1 into technical read 1 and biological read 1 in a `Python` version 3.10.13 environment using `dnaio` (Martin and Vorderman, 2023). Next, we filtered low-quality biological read 1 and read2 pairs by passing them as paired-end reads to `fastp` (Chen et al., 2018). In addition to the default setting, we disabled adapter trimming by specifying `disable adapter trimming`, because technical read 1s, instead of primer sequences, is adjacent to the 5′ end of biological read 1s. We specified that the required read length to pass the filtering is 10 bases by the `disable adapter trimming` argument.

### Read Alignment

The preprocessed biological read 1 and read2 pairs were aligned to the genome using `STAR` (Dobin et al., 2013) with an augmented gene annotation set. A *spliceu* STAR index was built for mouse and human using the 10X standard genome build and *spliceu* reference, respectively. To calculate the cDNA fragment length distribution, we then aligned biological read 1 and read2 pairs as paired-end reads using the *spliceu* STAR index. The alignments were exported as a BAM file and sorted by genome coordinate by passing `BAM SortedByCoordinate` to the `outSAMtype` argument. Additionally, a transcriptome BAM file was generated by passing `TranscriptomeSAM` to the `quantMode` parameter. We used the `Singleend` option of the `quantTranscriptomeBan` option to allow soft clips and gaps in transcriptome-based alignments. Additionally, we specified `outFilterScoreMinOverLread` and `outFilterMatchNminOverLread` as 0.33 to adjust this length-sensitive threshold to output the alignment for the split reads 1, that have much shorter sequence than read2, and set `alignSplicedMateMapLminOverLmate` as zero to allow more alignments in paired-end mapping.

Then, we used a similar setting to align just the read2s as single-end reads, and exported the alignments in the transcriptome-based coordinate. For accurate fragment length calculation, we filtered the genome-based alignments of the paired-end reads using `samtools` (Li et al., 2009) to ensure that the reads are properly and uniquely mapped to the genome by specifying `require-flags 2` and `d NH:1` for paired-end alignments. The genome-based paired-end alignments were used to calculate the cDNA fragment length and find the priming sites corresponding to each read (section 2.5). All types of alignments were used to build the validation read set(section 2.9).

### cDNA fragment length distribution

As introduced in section 2.3, we define the cDNA fragment length of a scRNA-seq read as the length of the cDNA insert of the corresponding cDNA library fragment. Given the genome alignment of a biological read 1, read2 pair, the cDNA fragment length of this read can be calculated as the contiguous genomic interval from the 5′-most locus to the 3′-most locus of the alignment. Only reads properly and uniquely mapped to the genome are used for calculating the cDNA fragment length distribution (section 2.5).

This process was done in a `Python` version 3.10.13 environment using `biopython` (Cock et al., 2009) and **pysam** ^‡^. We read the genome FASTA file using the `SeqIO` module from `biopython`. Next, given the genome alignment pair of a uniquely mapped paired-end read as an `AlignedSegment` (from `pysam`), we found the interval spanned by the paired-end read on the genome according to the corresponding `SAM` flags indicating their orientation and the `CIGAR` string. Specifically, we found the distance *D* between the closer ends of the paired-end alignment. Then, we sum the read length of biological read 1 and read 2, and *D* to get the final fragment length *L*. If the biological read 1 and read 2 intersect and they share gaps that are designated as *intronic* by `STAR` (as indicated by the character “N” in the `CIGAR` string), the gaps will not be included when calculating *D*. Additionally, the genomic sequence of the priming window captured by the oligo(dT) primer of this read, defined as the downstream 30-base genomic interval of biological read 1, was extracted from the genome according to the alignment. We also extracted a background genomic interval randomly selected around biological read 1. The extracted priming windows and background windows are used in section 2.7 for binding affinity prediction.

Note that the procedure is designed for processing reads generated from internal polyA priming that does not contain a gap (exon-exon junction) in the interval between the paired-end alignment. If the interval between a paired-end alignment, that is not covered by the reads themselves, spans an exon-exon junction, then the final fragment length will contain the length of the gap. If read 1 originates from polyA tail priming, then most likely, it will not align to the genome, and so the corresponding fragment will not be used in fragment length calculation (as intended), since it does not map as a proper pair. However, it is possible that, in some cases, read 1 overlaps the junction between the terminal exon and the polyA tail of a polyadenylated transcript. In such cases, the intergenic sequence downstream of the terminal exon in the genome will be regarded as the priming window, because the polyA tail of transcripts is not included in the genome build. However, our results suggested that these exceptional cases are rare in the selected datasets, as most fragment lengths fall into the expected range, from 190 to 290 base pairs, and most of the priming window sequences contain an A-SNR.

### Fragment length model fitting and evaluation

We fit the empirical scRNA-seq cDNA fragment lengths obtained from section 2.5 into a spline using `scipy` (Virtanen et al., 2020). As the expected fragment length ranges between 190 to 290 base pairs (section 2.3), we used all fragment lengths no larger than 1, 000 base pairs to fit the spline. For each training dataset, we calculated the frequency of all fragment lengths ranging from 1 to 1, 000 and normalized the frequencies to get the discrete empirical fragment length distribution. Then, we fit a spline on the distribution using `splrep` from `scipy` and set the smoothing condition argument, *s*, to 1/10^6^ to balance the fidelity and smoothness. The average and standard deviation of the root Mean Square Error (RMSE) of the prediction to the empirical distribution was calculated from the two test datasets.

The generality of the spline model was assessed using a 5-fold cross-validation experiment. In each iteration of the 5-fold cross-validation, we trained the model using eight training datasets and hold out the two validation sets. Then we calculated the predicted frequency from each model. The error bar plot showed the minimum and maximum difference between predictions of each model and the mean of the 5 models’ frequency prediction.

### The oligo(dT) binding affinity model

In section 2.5, we extracted the downstream sequence of biological read 1 corresponding to the empirical priming window together with background sequences. We then used these empirical priming window sequences and background sequences to train a multi-layer perceptron (MLP) using `MLPClassifier` from scikit-learn Pedregosa et al. (2011). By training an MLP on the priming windows, it should learn the sequence motifs present therein, and be able to predict the binding affinity of oligo(dT) primers for a given putative priming window.

We initialized the MLP using `MLPClassifier` with the `adam` optimizer. The maximum iterator parameter `max iter` was set to 500 because of the large size of the training data. As the extracted sequences might be intergenic instead of the actual priming window (in the minority of cases when the mapped read 1s originate from polyA tail priming), we first filtered out the sequences that do not contain an A-SNR of length at least 6 with at most one mismatch. Then, we trained the filtered priming window sequences from the eight training datasets in batches using `partial fit` because the training set is too large to load into memory at once. We held out 1, 000 within training examples from each dataset as the evaluation set. We then tested for overfitting of the trained MLP using the filtered priming windows from the two test datasets by comparing the mean accuracy between the evaluation and test sets. Note that the performance of the trained MLP might be limited by the potential mislabeled sequences in the provided training data. There are two sets of mislabeled sequences in our training data. The first set contains the background sequences that have the potential to be primed but were not selected for priming (false negatives). The second set contains the intergenic sequences extracted from the biological read 1 that are partially arising from polyA tails (false positives), because we extracted the upstream sequence of biological read 1s from the *genome*. However, Our results (not shown) suggests that the trained MLP can predict the holdout set and the test datasets consistently well, with a mean accuracy ∼ 80%.

### The Forseti model

In the evaluation of the splicing status of scRNA-seq reads, a critical piece of potentially useful evidence that is currently unused is the likelihood that particular priming sites give rise to the reads. For example, since the priming of sequenced fragments is expected to originate from the priming to A-SNRs along transcripts, and since fragment lengths in scRNA-seq follow a well-characterized distribution that can be inferred from observed data, one can evaluate the likelihood that a particular polyA tract gives rise to an observed read based on the mapping location of its read 2. Specifically, for a read *r* with a given mapping on a transcript *t*, starting at position *x*, one can consider this read to derive from the end of a cDNA fragment whose opposite end terminates proximate to a downstream A-SNR (i.e. the priming event associated with this fragment). To evaluate the probability of these potential cDNA fragments, one can evaluate the binding affinity of the A-SNR and the probability of observing a cDNA fragment of the implied length under the empirical fragment length distribution. If, for example, it is highly likely that a read is paired with an A-SNR located within an intron, then this provides strong evidence that the associated UMI should be assigned an unspliced status. On the other hand, if the read is likely paired with the poly-A tail, then the pairing is not particularly informative as to the associated UMI’s splicing status (since both spliced and unspliced molecules may be polyadenylated).

More formally, let the fragment length distribution be *f*. This is a probability distribution that assigns a probability *p*_*f*_ (*𝓁*) to observing a fragment of length *𝓁*. Let the binding affinity distribution *b* be another probability distribution that assigns a binding probability *p*_*b*_(*w*) for a potential priming window *w* containing an A-SNR based on its sequence. Further, for the start site *x* of a read mapping on exon *e* of transcript *t* of gene *g*, let pbs (*t, x*) = {(*w*_1_, *d*_1_), (*w*_2_, *d*_2_), … } denote the set of *all* potential priming windows downstream of *x* in *t* that contains an A-SNR, and for each such window *w*_*i*_ the corresponding distance *d*_*i*_ from *x*. Given a read *r* of ambiguous status that maps to position *x* on exon *e* of transcript *t*, we can evaluate the probability that this read arises from *t* (here, defined as the probability of the *most likely* associated binding event) as:

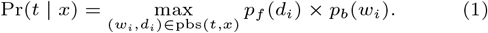

Then, assume that, in addition to *t*, the exon *e* belongs to other transcripts of the gene *g*. We define a function tx(*g, e, v*) to be a function that returns *all* transcripts of *g* that contains the exon *e* with the provided splicing status *v* ∈ {*s, u*}, where *s* stands for spliced and *u* stands for unspliced. Then, we can evaluate the probability that a read *r* arises from a spliced transcript of gene *g* as:

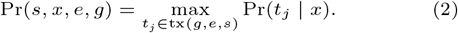

Finally, we define the probability that the read arises from a spliced transcript of gene *g* as:

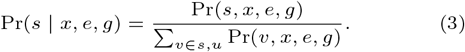

Additionally, if the read maps to multiple genes *G* = {*g*_1_, *g*_2_, … }, then the most appropriate gene *g* for explaining the read is defined as the gene that has the transcript yielding the maximum *P* (*v, x, e, g*) among all genes in *G*, i.e.:

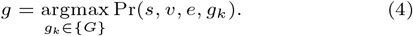

### Evaluation on experimental samples

In the experimental data on which we evaluated our model, we extracted sequencing reads for which read 2, when aligned alone, is of ambiguous splicing status, but when paired with biological read 1, the splicing status is determined (or almost certainly determined). In other words, we want to find the read pairs such that the read 2 is entirely contained within an exon but the biological read 1 is not entirely contained within the same exon. Specifically, biological read 1 could be intronic (i), span an intron-exon junction (i.e), span an exon-exon junction (e.e′), or be contained in a *different* exon than read 2 (e^∧^e′). In each of these cases, the definitive splicing status, either spliced or unspliced, determined from the biological read 1 served as the ground truth for the subsequent evaluation, where the model, along with the alignments of read 2 treated as single-end data, was used to predict the splicing status of the underlying fragment.

Here, using the alignments for paired-end reads from section 2.4, we briefly describe the process used to assign a definitive splicing status to reads that have an exonic read 2. We additionally note that reads that have pair-end alignments to multiple genes are ignored in the procedure (i.e. not considered for classification and subsequent evaluation) as we do not know the true gene of origin. First, we used `bedtools` (Quinlan and Hall, 2010) to mark all alignment positions for biological read 1 and read 2 that are entirely contained within exons according to the paired-end alignment records and the exon annotations generated in section 2.4. Again, if read 2 of a fragment is not contained within any exon, then read 2, by itself, is determinative as to the splicing status and the read pair will not be processed further. We retain the information about whether or not biological read 1 was entirely contained within an exon to aid in subsequent classifcation.

Given a biological read 1, read 2 pair, we obtain the reference transcripts explaining the paired-end read, denoted as {*T* ^pe^}, from its alignments. We also obtain the transcripts that are compatible with an *exonic* alignment of biological read 1 and read 2, denoted as 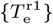 and 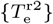, respectively.

We identify the inferred status of the read, as spliced or unspliced, as follows:

- (e.e′) reads: When (1) 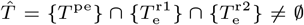 (2) 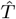 contains only spliced transcripts, (3) the corresponding exons in each 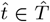 that are compatible with biological read 1 and read 2 are different, and (4) the genomic distance between the exons containing read 2 and biological read 1 in each 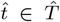 are *>* 1, 000 nucleotides apart, then we assign the read as a spliced (e.e′) read, because this means the fragment (but not either of the individual reads) spans an exon-exon junction of the common reference transcripts. Here, we require the distance between the involved exons to be larger than 1, 000 nucleotides to minimize mislabelling caused by short introns, where the fragment may place the reads on different exons and completely contain a short intervening intron.
- (e.e′)reads: When (1)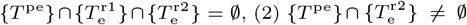 and (3) the intersection contains only spliced transcripts, we assign the read as a (e.e′) read, and therefore of spliced status, because this means its biological read 1 spans an exon-exon junction.
- i and i.e reads: When (1) 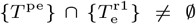 and (2) the intersection contains only unspliced transcripts, we assign the read as an unspliced read because this means its biological read 1 either crosses an intron-exon junction (i.e) or is complete contained within an intron (i), and therefore the read pair must have arisen from an unspliced molecule.

We processed each read with an exonic read 2 individually to get the spliced and unspliced labels for the evaluation set. We then applied our model using the alignments of read 2s, treated as single-end data, to predict the gene origin and the splicing status of the read. The model prediction of reads was compared with their assigned labels according to the above criteria to evaluate the performance of the model.

### Data Simulation

In addition to the experimental data, which employ a collection of rules involving the paired-end mapping of read 1 and read 2 to ascertain the true splicing status, we also evaluated our model on simulated data, where the true splicing status of each fragment is known with certainty. As with all simulations, the trade-off here is that the simulated data is, in general, simpler and “cleaner” than the experimental data. Nonetheless, as we observe similar AUCs in both cases (see Section 3), we believe that the simulated data analysis is a useful evaluation of `Forseti`.

We simulated paired-end reads from the *spliceu* transcriptomic reference(section 2.2). In order to mimic real data at both read-count and read-sequence levels, we seed our simulation with a count matrix from experimental data and introduced realistic sequencing errors into the simulated reads.

We applied `simpleaf` (He and Patro, 2023) to a 10X 1k Human PBMC v3 scRNA sample ^§^ using the *spliceu* reference to generate the spliced, unspliced and ambiguous count matrices. These matrices were loaded into Python as an AnnData object (Virshup et al., 2021) using `pyroe` He and Patro (2023).

As detailed in section 2.2, regions on the reference sequence with at least six consecutive adenine (A) bases are considered potential polyA priming sites. We denote by 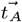the ordered list of potential polyA priming sites on transcript *t*. The polyA tail of each transcript was also considered a potential priming site.

Simulated reads were generated from the transcripts recorded in *spliceu* according to the following process. First, a gene *g* is selected for sequencing. For each gene *g* we draw *g*_*s*_ spliced reads and *g*_*u*_ unspliced reads, where these counts are determined by the corresponding counts from the processed sample. Given a gene *g* and a read status *v* ∈ {*u, s*}, we randomly select a transcript *t* of *g* with the desired splicing status *v*. Next, we select a polyA site from 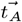 uniformly at random; let the site be denoted as 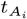 Then, we sampled a fragment length *d* according to the empirically derived fragment length distribution *f*. The pair of 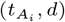 determines the underlying fragment being sampled, which spans the transcriptomic region of length *d* ending at 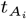. This determines the position of read 2 (the first 91 bases at the 5′) as well as the corresponding read 1 (the last 31 bases at the 3′), matching the cycle length of the reference dataset. The sequence of read 1 and read 2 were extracted from the spliceu reference sequences, which was loaded using the `SeqIO` module from `Biopython` (Cock et al., 2009).

As our focus in this work is on recovering the splicing status of fragments, the only simulated reads useful for our experiments are those where read 2 resides entirely within an exon. So as to not waste effort simulating and quantifying fragments where read 2 is, itself, of determined splicing status, we exclude fragments having a non-exonic read 2 from our simulated data. Thus, during simulation, we inspect each simulated potential fragment to determine if read 2 is ambiguous or not. If read 2 itself is determinative of the splicing status of the underlying molecule, we discard this read and sample a fragment again from the same gene and splicing status. To achieve this while still striving to obtain the desired read count for each gene, we implemented a resampling process, limited to a maximum of 100, 000 trials per read.

We note that, because we rejected and resampled “naturally simulated” fragments for which read 2 had a determinative splicing status, there is a selection effect among the fragments that eventually complete the simulation process, and that the simulated data is therefore not entirely random with respect to the underlying collection of polyA sites and the target fragment length distribution. Nonetheless, we find that this process performs reasonably well in terms of generating fragments whose ambiguity profiles match those of experimentally ambiguous fragments.

Moreover, it is also important to note that two specific categories of reads are *fundamentally ambiguous* and remain indistinguishable within our model and, in fact, in the context of any model that just considers the type of information used in `Forseti`. The first category includes fragments where read 1.is from the polyA tail and read 2 resides in the terminal exon of the transcript of origin. Such fragments can originate from either the spliced or unspliced versions of a transcript, leading to inherent and fundamental ambiguity. The second category comprises read pairs where the whole fragment (read 2.and its mate read 1) are from *the same exon* within a transcript. These are *fundamentally ambiguous* for the same reason as the first category but tend to be rare in practice as typical fragment lengths are longer than typical exon lengths. To test the performance of our model with and without fundamental ambiguity, on top of the data simulated by the procedure described above, we also simulated a read set without fundamental ambiguity. In this set, we require not only that read 2 of a fragment is contained in an exon, but also that its corresponding read 1 is not contained within the same exon, and the fragment is not generated from the polyA tail of transcripts.

To mimic realistic sequencing error profiles, we employed `InSilicoSeq` (`ISS`) (Gourlé et al., 2019) to introduce realistic Illumina errors into the simulated reads. We constructed a custom error model using the reference 10X 1k PBMC sample data mentioned above. We aligned the data with `STAR` using the *spliceu* annotation. `STAR` was executed with similar flags as in section 2.4 for aligning only read 2, but with an additional flag to include the `MD` tag in the output BAM file. The error model was then built using `ISS` with the default setting. `ISS` estimated the quality score for each base, thereby introducing substitution errors into the simulated reads based on the estimated scores. Finally, we exported the read pairs to two separate `FASTQ` files using `SeqIO` from `Biopython`.

## Results

In this study, we developed a probabilistic model, `Forseti`, to infer the splicing status of the molecule origin for scRNA-seq reads. Our model makes use of a fragment length distribution and a binding affinity model learned from empirical data to identify and score the potential sequenced fragments that led to the observed scRNA-seq reads. From these scored fragments, `Forseti` can not only predict the splicing status of the molecule origin that the read was drawn, but also recognize the true gene origin when reads are compatible with multiple genes. We evaluated `Forseti` on both simulated and experimental validation sets, and found that `Forseti` precisely predicts the splicing status of reads in all validation datasets, with AUC scores ranging from 0.85 to 0.93 and an average AUC of 0.90.

### Evaluation of model components

In most tagged-end scRNA-seq protocols, the cDNA fragments are generated with a preferred length of 300-400 base pairs, which means the cDNA fragment length in scRNA-seq might follow a distribution. However, the potential cDNA distribution is challenging to model, or even detect, in most existing scRNA-seq data, because the recommended sequencing format captures only the 3′ end sequence of the cDNA insert. To overcome this challenge and model the cDNA fragment length distribution, we collected ten scRNA-seq datasets that were generated by applying the alternative sequencing format (10x Genomics, 2018), in which read 1 is sequenced the same cycles as read 2, so that the sequence of the 5′ *and* 3′ end of the cDNA insert in each sequenced fragment are both captured, by read 1 and read 2, respectively. In other words, these data can be treated as paired-end data, and the cDNA fragment length of each sequenced fragment can be calculated from the alignments of the paired-end reads. In this work, we fit a cubit spline model on the empirical fragment lengths calculated according to the paired-end read alignments from eight training datasets selected from the collected datasets. We evaluated its performance on the two holdout testing datasets (section 2.5). As shown in fig. 2a, the distribution model’s close fit to data, strikes a balance between smoothness and fidelity. The low Root Mean Square Error (RMSE) score in our evaluation indicates a high accuracy of the model in predicting the frequency of fragment length within two test datasets(RMSE mean = 3 × *e*^−4^, RMSE standard deviation = 1 × *e*^−4^). In addition, we assessed the trained spline model’s generalizability with a 5-fold cross-validation experiment. For each fold, we trained a cubic spline model with eight training datasets. The error bar plot in fig. 2b illustrates that minimal variation is observed in the frequency prediction across all five folds, indicating model’s consistent performance and a generally similar fragment length distribution across samples. This consistency also supports our initial hypothesis regarding a generalized insert size distribution and that a distribution can be learned and robustly applied to various datasets across different species, cycle length and chemistry version.

**Fig. 1.**
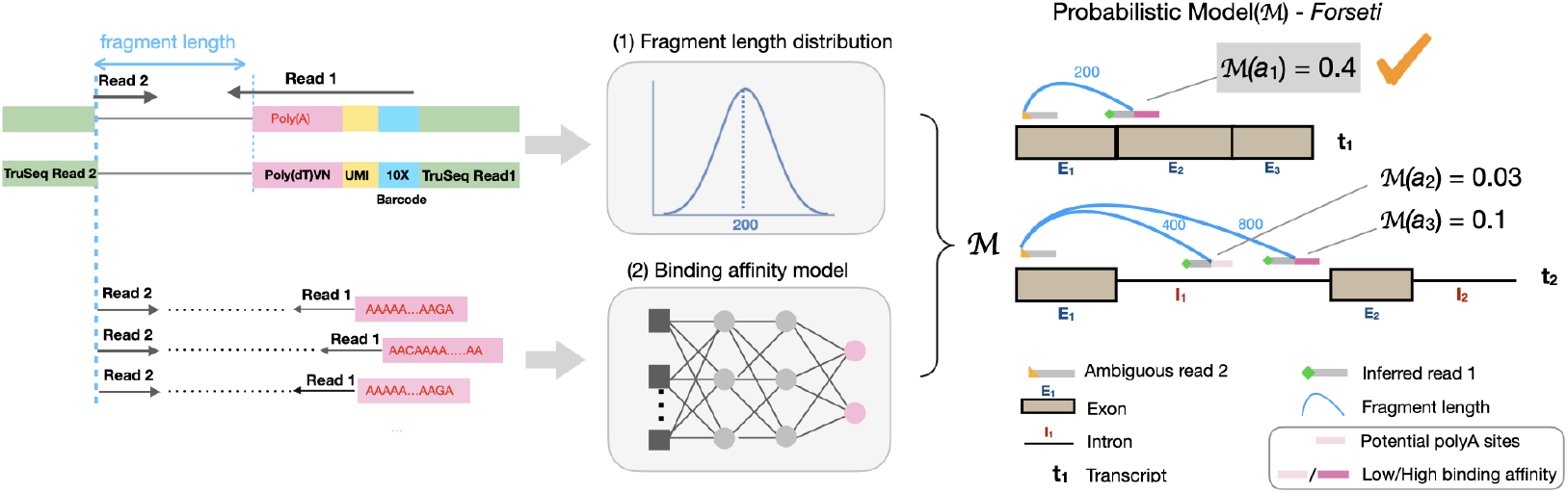
Overview of the `Forseti` model of splicing status inference. The fragment length and binding affinity models are used to score putative fragments, and splicing status of the molecule giving rise to the highest-scoring fragment yields the prediction.

**Fig. 2.**
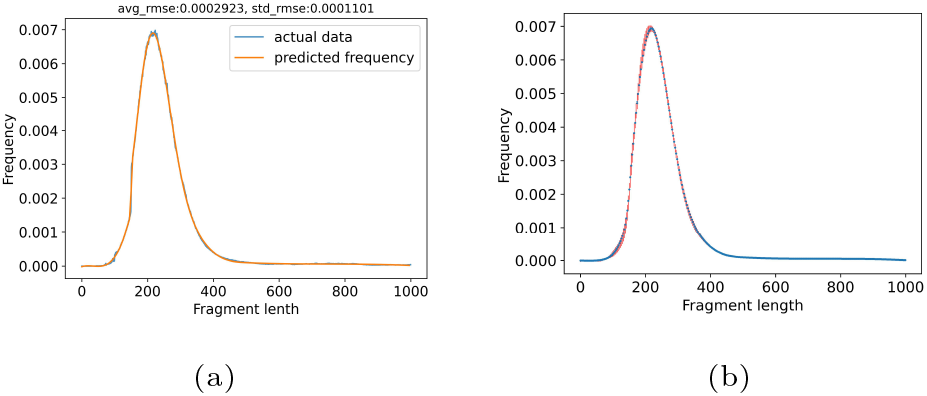
We fit a generalized distribution model of fragment length frequency. The accuracy was assessed by Root Mean Square Error (RMSE) score on test datasets(a).The blue line is the mean of the frequency from 2 test datasets, and the orange line is the predicted frequency from the trained spline model. The consistency of model across samples was evaluated with a 5-fold cross-validation was showed in the Error Bar plot (b).

In addition to the fragment length model, `Forseti` utilizes a multi-layer perceptron (MLP) to predict the binding affinity of the potential priming window containing the sequence motif where the poly(T) primer binds. The MLP was trained and tested on experimental priming windows obtained from the training and test split used in the fragment length model. The mean accuracy of the trained MLP on the training datasets, the holdout set from the training data, and the tested datasets is 0.877, 0.865, and 0.860, respectively, suggesting the high accuracy and generalizability of the trained MLP for identifying the experimental priming windows from random background sequences across all datasets.

### Forseti accurately predicts the splicing status of reads

We evaluated `Forseti` on four evaluation read sets, two from experimental data, and the other two from simulated data (sections 2.9 and 2.10). Because `Forseti`, and in fact, any model that just uses the same information, is unable to predict a definitive splicing status for *fundamentally ambiguous* fragments — fragments arising either from polyA tail priming and staying in the terminal exon, or having both biological read 1 and read 2 entirely contained within the same exon — we generated two sets of simulated data; one with and one without fundamental ambiguity (section 2.10).

The model performance was assessed using the receiver operating characteristic (ROC) curve and, specifically, by evaluating the area under ROC curve (AUC) scores. The ROC curve plots the true positive rate and false positive rate from all predictions as the classification threshold is swept across the value of all scores. The AUC, ranging from 0 to 1, measures the area under the ROC curve. A high AUC score means a model has a good prediction power — that the probability assigned by our model to a splicing status is well-correlated with the true splicing status. As shown in fig. 3, our model demonstrated consistently high AUC among all evaluated read sets. In particular, the AUC of `Forseti` on the two experimental sets generated from human and mouse scRNA-seq datasets is 0.89 and 0.85 (figs. 3c and 3d), suggesting the generality of our model across different species and cell types. The high AUC scores of `Forseti` on the simulated data with and without fundamental ambiguity (figs. 3a and 3b), consistent with the experimental sets, suggest that the mechanism — internal polyA priming — we used to generate the simulated dataset (section 2.10) well-mimics the real-world mechanism that generates ambiguous reads. Also, our result highlights that including fundamental ambiguity did not hamper the ability of our model to predict the correct splicing status for reads when possible. We also included the ROC curve of three baseline models, implying the current best practices for resolving splicing ambiguity (La Manno et al., 2018; He et al., 2022; 10x Genomics, 2022b; Kaminow et al., 2021; Eldjárn Hjörleifsson et al., 2022b) Figure 3. These baseline methods either assign all reads a spliced status probability of 1 (“All spliced”), 0 (“All unspliced”), or a random probability ranging from 0 to 1 (“All random”). This “All random” predictor is generated by first assigning a random probability in [0, 0.5] to each truly unspliced read and a random probability selected in [0.5, 1] to each spliced read, and then randomly shuffling the order of these probabilities. All three baseline models had an AUC=0.5, suggesting that no trivial predictor can extract meaningful splicing status at a level approaching that achieved by `Forseti`.

**Fig. 3.**
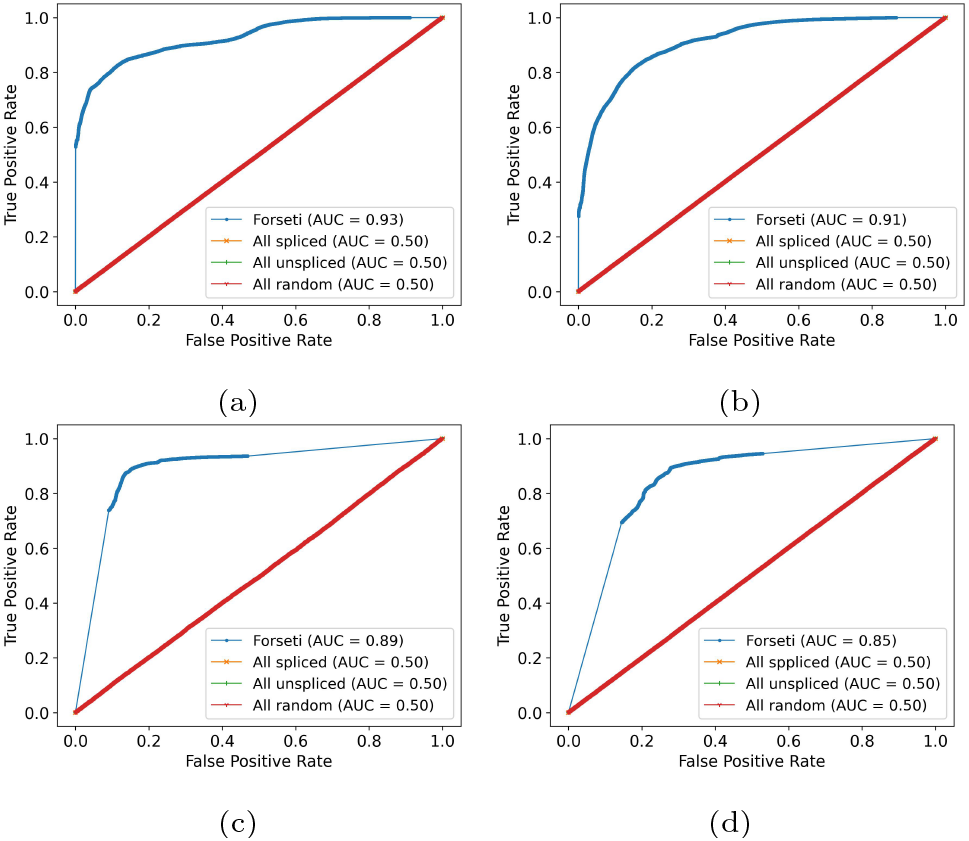
`Forseti` accurately predicts the splicing status of both simulated and experimental reads, evaluated using receiver operating characteristic (ROC) curve and Area Under ROC (AUC) score. In all plots, the blue curve represents the ROC of `Forseti`, and the other three curves stacking with each other, represent the three baseline models. Panel (a) and (b) show the ROC curve and the corresponding Area Under ROC (AUC) of simulated with and without fragments entirely contained within an exon and fragments arising from terminal polyA priming, respectively. Panel (c) and (d) show the AUC of the two experimental datasets from human and mouse, respectively.

One advantage of being a probabilistic model is that, for those fundamentally ambiguous reads that our model is currently unable to handle, their splicing status will remain ambiguous in `Forseti`’s prediction, with a probability of being spliced as exactly 0.5. In the two experimental datasets, reads that remain with an ambiguous splicing status after `Forseti` account for 14% and 19%, respectively, suggesting that fundamental ambiguity is cell-type specific. In the simulated data with fundamental ambiguity, ambiguous reads account for more than 76% of reads, mainly because our simulation setting prefers generating fundamentally ambiguous reads (section 2.10). In the simulated dataset without fundamental ambiguity, this percent becomes 15%, coincident with the experimental sets. This suggests that our simulation that allows fundamental ambiguity is unrealistic (i.e. is overly pessimistic) and that the degree of such ambiguity is likely substantially lower in real-world, experimental data. In fig. 4, we show similar ROC plots for the evaluation sets but with ambiguous predictions included. Compared with figs. 3a and 4a, the AUC of these rather unrealistic simulations, in which fundamentally ambiguous reads are dominated, decreased from 0.93 to 0.68. Though a large decrease from 0.93, this is still much higher than the 0.5 AUC obtained by the baseline models. Apart from this overly-pessimistic case, the AUC of other evaluation sets remains consistently high. We note that the middle part of the curve, not covered by examples (i.e. the thinner part of the line in fig. 4), is caused by ambiguous predictions, as `Forseti` assigned a probability of 0.5 to all of them.

**Fig. 4.**
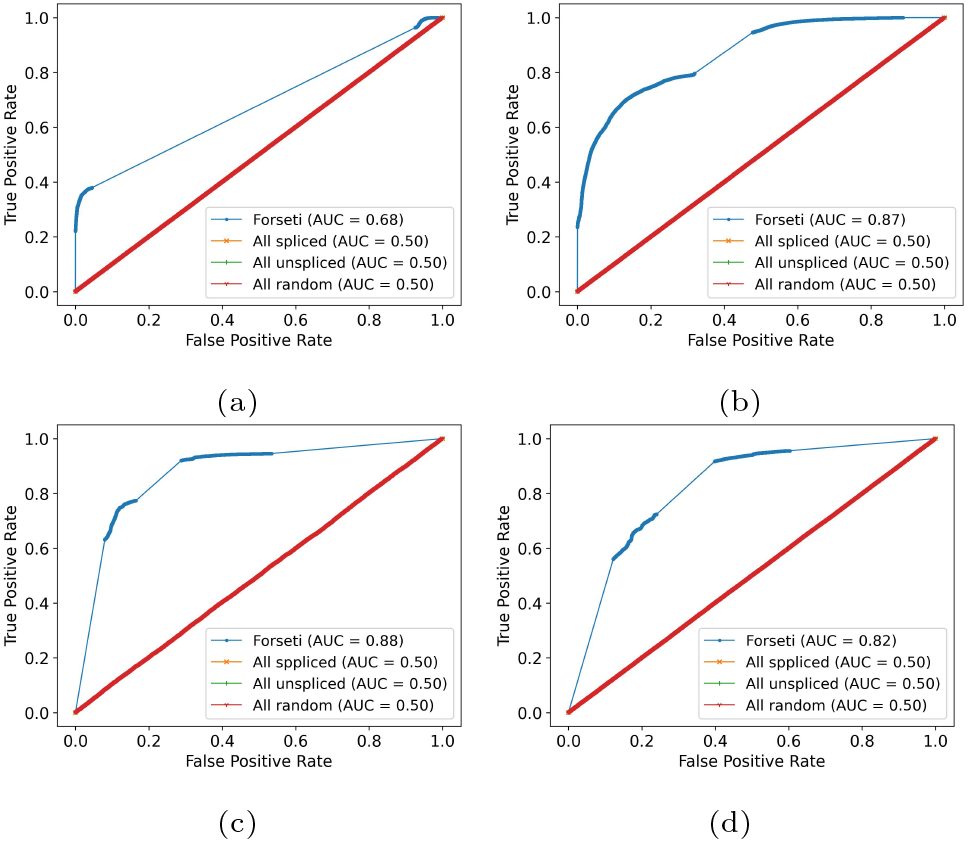
This figure shows the ROC curves that are analogous to those in fig. 3, except where reads predicted as having an ambiguous splicing status have been included (section 3).

### Forseti can disambiguate gene multimapping reads

Another advantage of being a probabilistic model is that the predicted probabilities of `Forseti` can be regarded as the confidence score that each potential downstream priming site, when paired with an alignment for read 2, represents that actual underlying fragment from which read 2 was sequenced. The final splicing status of a read with respect to a gene is assigned as the splicing status of the reference transcript for which the most confident alignment, priming site pair occurs. Similarly, this highest confidence score (probability) can *also* be used to evaluate the true gene origin of reads compatible with multiple genes (which may arise because of overlapping genes in the genome, or because of sequence-similar genes, such as members of a gene family).

To select a splicing status among spliced and unspliced transcripts within a gene, we simply compare the highest scoring fragment from any spliced transcript to the highest scoring fragment from any unspliced transcript. However, the same underlying predictive scoring mechanism of `Forseti` can also be used to evaluate the relative probability of the truly generating fragment *between* genes.

Specifically, consider a read that is multimapping between two genes *g*_1_ and *g*_2_. For each gene, `Forseti` computes a score for all potential fragments and records the highest score among all transcripts of *g*_1_ and all transcripts of *g*_2_, respectively. Then, we can then compare the highest score computed for *g*_1_ and *g*_2_ to determine the best gene origin of the read between these genes. In the current work, we consider the maximum score as the criterion when comparing compatible genes. However, we note that other aggregation mechanisms, such as the sum, the expected fragment probability, or a weighted expectation, can also be used for scoring genes, and evaluating optimal aggregation mechanisms is an interesting direction for future work.

One finding from our result is that, in our simulated data, multi-gene mapped reads were all simulated from unspliced transcripts, mainly from the regions that are shared by multiple genes. Because the sequence of these shared regions is identical, the true gene origin of these reads is indistinguishable in our model. However, for the two experimental evaluation datasets, `Forseti` assigned a single best gene origin to 65% of the multigene mapped reads in one dataset and to 93% of the multi-gene mapped reads in the other (the remaining reads had a tied highest score among genes, and thus no single gene prediction could be made). However, when a single gene prediction could be made, `Forseti`’s prediction was correct in 74% and 63% of the cases, respectively (i.e., the true gene origin of the multi-gene mapped reads was identified). This suggests that in reality, most multi-gene mapped reads are distinguishable, and, among the distinguishable subset of multi-gene mapped reads, `Forseti` can assign the true gene origin to them in the majority of cases. As the gene origin assignment is affected by the weight `Forseti` places on internal polyA compared to polyA tail priming, a weight parameter naturally arises that affects the relative preference for polyA tails versus internal polyA priming. Exploring the trade-off between the accuracy of gene-origin prediction and splicing status prediction by optimizing or altering this weight is an interesting direction for future work. Here, however, we consider just the simple case where we treat the two polyA classes equally.

While the rescue of multi-gene reads is not the primary focus of our model, and was not a target application during `Forseti`’s design, the fact that it provides meaningful predictions in this distinct task is evidence of the promise that fundamentally better fragment modeling can make to scRNA-seq pre-processing. To our knowledge, our model is the first to attempt to resolve fragment-level multi-gene mapping in scRNA-seq data, thanks to the utilization of the fragment length distribution and binding affinity models. Further, these informative predictions stack on top of existing models for partially allocating gene-multi-mapping UMIs, such as the EM approach introduced in Srivastava et al. (2019) and later adopted in Melsted et al. (2021); Kaminow et al. (2021) and He et al. (2022).

## Conclusion

In this work, we introduce `Forseti`, a mechanistic and predictive model of the splicing status of the sequenced fragments in scRNA-seq samples. As the first predictive model of which we are aware that focuses on resolving the splicing status ambiguity in scRNA-seq, `Forseti` utilizes the fragment length distribution and a binding affinity model trained on a variety of experimental scRNA-seq datasets to predict the probability that the observed read is associated with specific sequenced fragments arising from each compatible reference transcript. Our results show that the AUC of our model on both experimental and simulated evaluation sets is high, demonstrating the consistent performance of `Forseti` across species and cell types. Furthermore, by virtue of being a probabilistic model that seeks to score potential fragments of origin of a read, and not *just* the read’s splicing status, our model predictions can also be compared across genes and used to help successfully resolve the gene of origin for the majority of multi-gene mapped reads in our experimental evaluation datasets.

Because `Forseti` predicts the splicing status of sequenced fragments based on the difference of the nucleotide sequence of the spliced and unspliced transcripts, one limitation of `Forseti` is that, when the possible fragments in spliced and unspliced transcripts are identical (for example for fragments are entirely contained within an exon) `Forseti` will be unable to resolve the splicing status of the fragments. However, one potential path to overcoming this limitation is to aggregate evidence over all fragments from the same unique molecule identifier. In this case, an individual read may be *fundamentally ambiguous*, but other fragments of the same UMI may have a definitive splicing status or at least an informative splicing probability, so as to be used to distinguish the splicing status of their common molecule of origin. Likewise, another potential use case for `Forseti` at the UMI-level is to aid in determining potential UMI collisions. If a UMI contains conflicting evidence, i.e., some of its sequenced fragments are predicted as spliced and others as unspliced, this might indicate UMI collision, suggesting the potential of using `Forseti` to also improve the UMI resolution in scRNA-seq. Finally, we have developed `Forseti` as a proof-of-concept model, demonstrating the potential and promise of fragment-level modeling to elucidate splicing status in scRNA-seq data. However, one important direction for future work is to properly integrate this model into existing tools for efficient scRNA-seq processing, like our `alevin-fry` tool He et al. (2022). While not conceptually difficult, such integration will require the propagation of additional information through the processing pipeline, and it will also likely highlight the need and opportunity for further computational enhancements and simplifications that will make the fragment probabilities evaluated in `Forseti` faster to calculate at the scale of ever-growing scRNA-seq experiments.

## Author contributions statement

R.P. supervised the work. D.H. and R.P. designed the model. D.H., Y.G., and S.C. generated the fragment length distribution. D.H., N.Q., Y.G., and R.P., designed the classification rules used for fragment assignment, and D.H., Y.G. and N.Q. worked on implementations of these rules. D.H. trained the multi-layer perception. Y.G. fit the spline model with the help of D.H. and R.P.. D.H. developed the Nextflow workflow with the help of Y.G.. D.H. and Y.G. generated the experimental and simulated evaluation data with the help of R.P.. D.H. implemented `Forseti` with the help of Y.G. and R.P.. D.H., Y.G., and R.P. wrote the manuscript. All authors reviewed the manuscript. N.Q. carried out this work as a participant in the BRIDGE REU program at the University of Maryland.

## Acknowledgments

This work has been supported by the US National Institutes of Health (R01 HG009937), and the US National Science Foundation (CCF-1750472, and CNS-1763680). Also, this project has been made possible in part by grant number 252586 from the Chan Zuckerberg Initiative Foundation. The founders had no role in the design of the method, data analysis, decision to publish, or preparation of the manuscript. We thank Dr. Stephen M. Mount and Dr. Najib M. El-Sayed from the University of Maryland, College Park for discussing the idea of exploring paired-end scRNA-seq data. We thank Mervin Fansler from the Memorial Sloan Kettering Cancer Center for providing us with a curated collection of metadata for publicly available paired-end scRNA-seq datasets.

## Footnotes

* https://kb.10xgenomics.com/hc/en-us/articles/360000939852-What-is-the-difference-between-Single-Cell-3-and-5-Gene-Expression-libraries-

† https://support.10xgenomics.com/single-cell-gene-expression/software/downloads/7.0/

‡ https://github.com/pysam-developers/pysam

§ https://www.10xgenomics.com/datasets/1-k-pbm-cs-from-a-healthy-donor-v-3-chemistry-3-standard-3-0-0

## References

10x Genomics (2018). Technical Note – Base Composition of Sequencing Reads of Chromium Single Cell 3’ v2 Libraries, Document Number CG000080, 10x Genomics, (2018, November 19).

10x Genomics (2021). Technical Note – Interpreting Intronic and Antisense Reads in 10x Genomics Single Cell Gene Expression Data, Document Number CG000376, 10x Genomics, (2021, August 9).

10x Genomics (2022a). Technical Note – Assay Scheme and Configuration of Chromium Single Cell 3’ v2 Libraries, Document Number CG000108, 10x Genomics, (2022, December 2).

10x Genomics (2022b). Technical Note – Interpreting Single Cell Gene Expression Data With and Without Intronic Reads, Document Number CG000554, 10x Genomics, (2022, June 21).

Bergen, V., M. Lange, S. Peidli, F. A. Wolf, and F. J. Theis (2020, August). Generalizing rna velocity to transient cell states through dynamical modeling. Nature Biotechnology 38 (12), 1408–1414.

Chamberlin, J. T., Y. Lee, G. T. Marth, and A. R. Quinlan (2022, August). Differences in molecular sampling and data processing explain variation among single-cell and singlenucleus RNA-seq experiments.

Chen, S., Y. Zhou, Y. Chen, and J. Gu (2018, September). fastp: an ultra-fast all-in-one fastq preprocessor. Bioinformatics 34 (17), i884–i890.

Chen, X., P. Roelli, D. Hereñú, P. Höjer, and T. Stuart (2023, October). Teichlab/scg lib structs: Release 26th Oct 2023.

Cock, P. J. A., T. Antao, J. T. Chang, B. A. Chapman, C. J. Cox, A. Dalke, I. Friedberg, T. Hamelryck, F. Kauff, B. Wilczynski, and M. J. L. de Hoon (2009, March). Biopython: freely available python tools for computational molecular biology and bioinformatics. Bioinformatics 25 (11), 1422–1423.

Dobin, A., C. A. Davis, F. Schlesinger, J. Drenkow, C. Zaleski, S. Jha, P. Batut, M. Chaisson, and T. R. Gingeras (2013). Star: ultrafast universal rna-seq aligner. Bioinformatics 29 (1), 15–21.

Eldjárn Hjörleifsson, K., D. K. Sullivan, G. Holley, P. Melsted, and L. Pachter (2022a, December). Accurate quantification of single-nucleus and single-cell rna-seq transcripts.

Eldjárn Hjörleifsson, K., D. K. Sullivan, G. Holley, P. Melsted, and L. Pachter (2022b, December). Accurate quantification of single-nucleus and single-cell rna-seq transcripts.

Gorin, G., S. Yoshida, and L. Pachter (2023, October). Assessing markovian and delay models for single-nucleus RNA sequencing. Bulletin of Mathematical Biology 85 (11).

Gourlé, H., O. Karlsson-Lindsjö, J. Hayer, and E. Bongcam-Rudloff (2019). Simulating illumina metagenomic data with insilicoseq. Bioinformatics 35 (3), 521–522.

He, D. and R. Patro (2023, March). simpleaf: A simple, flexible, and scalable framework for single-cell transcriptomics data processing using alevin-fry.

He, D., C. Soneson, and R. Patro (2023, January). Understanding and evaluating ambiguity in single-cell and single-nucleus RNA-sequencing.

He, D., M. Zakeri, H. Sarkar, C. Soneson, A. Srivastava, and R. Patro (2022, March). Alevin-fry unlocks rapid, accurate and memory-frugal quantification of single-cell RNA-seq data. Nature Methods 19 (3), 316–322.

Kaminow, B., D. Yunusov, and A. Dobin (2021). STARsolo: accurate, fast and versatile mapping/quantification of single-cell and single-nucleus RNA-seq data. BioRxiv.

La Manno, G., R. Soldatov, A. Zeisel, E. Braun, H. Hochgerner, V. Petukhov, K. Lidschreiber, M. E. Kastriti, P. Lönnerberg, A. Furlan, J. Fan, L. E. Borm, Z. Liu, D. van Bruggen, J. Guo, X. He, R. Barker, E. Sundström, G. Castelo-Branco, P. Cramer, I. Adameyko, S. Linnarsson, and P. V. Kharchenko (2018, August). RNA velocity of single cells. Nature 560 (7719), 494–498.

Lawrence, M., R. Gentleman, and V. Carey (2009, May). rtracklayer: an r package for interfacing with genome browsers. Bioinformatics 25 (14), 1841–1842.

Lawrence, M., W. Huber, H. Pages, P. Aboyoun, M. Carlson, R. Gentleman, M. Morgan, and V. Carey (2013). Software for computing and annotating genomic ranges. PLoS Computational Biology 9.

Li, H., B. Handsaker, A. Wysoker, T. Fennell, J. Ruan, N. Homer, G. Marth, G. Abecasis, and R. D. and (2009, June). The sequence alignment/map format and SAMtools. Bioinformatics 25 (16), 2078–2079.

Li, S., P. Zhang, W. Chen, L. Ye, K. W. Brannan, N.-T. Le, J.-i. Abe, J. P. Cooke, and G. Wang (2023, April). A relay velocity model infers cell-dependent rna velocity. Nature Biotechnology.

Martin, M. and R. H. P. Vorderman (2023). dnaio: Efficiently read and write sequencing data from python.

Melsted, P., A. S. Booeshaghi, L. Liu, F. Gao, L. Lu, K. H. J. Min, E. da Veiga Beltrame, K. E. Hjörleifsson, J. Gehring, and L. Pachter (2021). Modular, efficient and constant-memory single-cell RNA-seq preprocessing. Nature Biotechnology, 1–6.

Nam, D. K., S. Lee, G. Zhou, X. Cao, C. Wang, T. Clark, J. Chen, J. D. Rowley, and S. M. Wang (2002, April). Oligo(dT) primer generates a high frequency of truncated cDNAs through internal poly(a) priming during reverse transcription. Proceedings of the National Academy of Sciences 99 (9), 6152–6156.

Pages, H. (2023). BSgenome: Software infrastructure for efficient representation of full genomes and their SNPs. R package version 1.68.0.

Pages, H., P. Aboyoun, R. Gentleman, and S. DebRoy (2023). Biostrings: Efficient manipulation of biological strings. R package version 2.68.1.

Pedregosa, F., G. Varoquaux, A. Gramfort, V. Michel, B. Thirion, O. Grisel, M. Blondel, P. Prettenhofer, R. Weiss, V. Dubourg, J. Vanderplas, A. Passos, D. Cournapeau, M. Brucher, M. Perrot, and E. Duchesnay (2011). Scikit-learn: Machine learning in Python. Journal of Machine Learning Research 12, 2825–2830.

Pool, A.-H., H. Poldsam, S. Chen, M. Thomson, and Y. Oka (2023, September). Recovery of missing single-cell RNA-sequencing data with optimized transcriptomic references. Nature Methods 20 (10), 1506–1515.

Quinlan, A. R. and I. M. Hall (2010, January). BEDTools: a flexible suite of utilities for comparing genomic features. Bioinformatics 26 (6), 841–842.

Soneson, C., A. Srivastava, R. Patro, and M. B. Stadler (2021, January). Preprocessing choices affect rna velocity results for droplet scrna-seq data. PLOS Computational Biology 17 (1), e1008585.

Srivastava, A., L. Malik, T. Smith, I. Sudbery, and R. Patro (2019). Alevin efficiently estimates accurate gene abundances from dscRNA-seq data. Genome Biology 20 (1), 1–16.

Stark, R., M. Grzelak, and J. Hadfield (2019, July). RNA sequencing: the teenage years. Nature Reviews Genetics 20 (11), 631–656.

Svoboda, M., H. R. Frost, and G. Bosco (2022, March). Internal oligo(dT) priming introduces systematic bias in bulk and single-cell RNA sequencing count data. NAR Genomics and Bioinformatics 4 (2).

Virshup, I., S. Rybakov, F. J. Theis, P. Angerer, and F. A. Wolf (2021, December). anndata: Annotated data.

Virtanen, P., R. Gommers, T. E. Oliphant, M. Haberland, T. Reddy, D. Cournapeau, E. Burovski, P. Peterson, W. Weckesser, J. Bright, S. J. van der Walt, M. Brett, J. Wilson, K. J. Millman, N. Mayorov, A. R. J. Nelson, E. Jones, R. Kern, E. Larson, C. J. Carey, İ. Polat, Y. Feng, E. W. Moore, J. VanderPlas, D. Laxalde, J. Perktold, R. Cimrman, I. Henriksen, E. A. Quintero, C. R. Harris, A. M. Archibald, A. H. Ribeiro, F. Pedregosa, P. van Mulbregt, and SciPy 1.0 Contributors (2020). SciPy 1.0: Fundamental Algorithms for Scientific Computing in Python. Nature Methods 17, 261–272.

